# Designing human Sphingosine-1-phosphate lyases using a temporal Dirichlet variational autoencoder

**DOI:** 10.1101/2022.02.14.480330

**Authors:** Evgenii Lobzaev, Michael A. Herrera, Dominic J. Campopiano, Giovanni Stracquadanio

## Abstract

Enzymatic deficiencies cause the accumulation of toxic levels of substrates in a cell and are associated with life-threatening pathologies. Restoring physiological enzymes levels by injecting a recombinant version of the defective enzyme could provide a viable therapeutic option. However, these enzyme replacement therapies have had limited success, as the recombinant enzymes are less catalytically active, cause immune response and are difficult to manufacture. Moreover, the vast sequence design space makes finding enzymes with desired therapeutic properties extremely challenging.

Here, we present a new enzyme engineering framework, which builds on recent advances in deep learning, variational calculus and natural language processing, to design variants of human enzymes with biochemical features comparable to the wild type protein as a way to rapidly build targeted libraries for downstream screening. We applied our method to design variants of human Sphyngosine-1-phosphate lyase (HsS1PL) as potential therapeutic treatments for nephrotic syndrome type 14 (NPHS14), and characterized their biochemical properties through extensive sequence and molecular dynamics analyses.

## Introduction

Sphingolipids (SL), and their N-acylated derivatives, ceramides, are ubiquitous components of eukaryotic cell membranes where they play essential structural roles [1]. However, in recent years evidence continues to grow that they are also important players in various other pathways, such as cell signalling, survival and regulation [1, 2]. Moreover, studies of wide-ranging diseases, such as Type II Diabetes (T2D), Alzheimer’s, inflammatory diseases [3], Parkinson’s [4] and early onset Amyotrophic lateral sclerosis (ALS, [5]) are beginning to link sphingolipid and ceramides in numerous pathologies [6].

Within the complex lipidomic inventory, sphingosine 1-phosphate (S1P) is a key molecule whose metabolism is closely regulated [7]. An important enzyme that degrades S1P is the pyridoxal 5’-phosphate (PLP)-dependent S1P lyase (S1PL) [8, 9]. It was shown that S1PL catalyses the conversion of S1P to phospho-ethanolamine and (2E)-hexadecenal and both these products can be recycled through various metabolic pathways. Therefore, S1PL is a key regulatory node to control cellular S1P levels and metabolic flux through the pathway. S1PL has also been linked to many diseases, including rare S1PL insufficiency syndromes [10]; in particular, autosomal recessive loss of function mutations at the *SGPL1* locus encoding the S1PL is associated with nephrotic syndrome type 14 (NPHS14), which causes progressive renal dysfunction and leads to kidney failure [11, 12, 13]. Currently, no treatment is available for NPHS14, but S1P metabolism has been proposed as a potential therapeutic target; in particular, restoring S1PL function by injecting a recombinant version of the defective enzyme could prove useful, similar to existing enzyme replacement therapies for sphingolipid related pathologies such as Gaucher’s and Fabry’s disease [14].

However, designing effective therapeutic enzymes has been challenging; in particular, synthetic enzymes have had limited success [15], as they have poor catalytic activity, they are unstable in blood, can cause immune response, and are difficult to deliver at a sustainable cost at the point of care [16].

Identifying effective therapeutic enzymes could be possible by efficiently building targeted libraries of variants that can be characterized downstream to select those with the desired therapeutic properties [17]. Variants of wild type enzymes can be identified using molecular approaches, such as Directed Evolution (DE) [18], but they are usually expensive and low throughput, as they are limited by the ability to rapidly build and screen variants at scale [19]. Computational approaches, instead, have been instrumental in streamlining protein design by several orders of magnitude [20]. However, current methods are usually limited by the number of homolog sequences and the corresponding quality of multiple sequence alignments, along with the availability of known tertiary structures. Recently, instead, deep generative learning has proven to be a viable solution to generate new, unobserved functional proteins [21], either by learning evolutionary constraints from highly curated multiple sequence alignments [22] or directly from protein sequences [23, 24]. However, these methods require a large number of sequences to be effectively trained and extensive computational resources.

Here we addressed these problems by developing a deep generative model, called Temporal Dirichlet Variational Autoencoder (TDVAE), which allows to encapsulate enzyme features extracted directly from their primary structure found across species into a low dimensional discrete-like, multimodal statistical space, that can be efficiently explored to generate variant enzymes with wild type like biochemical properties.

We then used TDVAE to design a library of variants of the human Sphyngosine-1-phosphate lyase (HsS1PL) and showed, through extensive sequence and molecular dynamics analyses, that they retain wild type biochemical properties and are viable candidates for downstream experimental testing.

## Methods

### A deep generative learning framework for enzyme engineering

Enzyme engineering requires learning how to sample the protein design space to identify amino acid sequences associated with a desired catalytic function. Here we hypothesize that the design space has a statistical structure, whose functional form and corresponding parameters are unknown but can be learned from known enzyme sequences.

We hereby assume that the probability of observing an enzyme sequence *x* depends on a latent random variable *z*, such that *p*_*θ*_(*x, z*) = *p*_*θ*_(*x*|*z*)*p*_*θ*_(*z*), with *p*_*θ*_(*x*|*z*) and *p*_*θ*_(*z*) being parametric distributions. Thus, an enzyme sequence can be considered the result of a generative process, which involves sampling a random variable 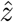 from *p*_*θ*_(*z*), and then building a sequence 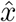 by sampling from the conditional probability 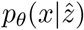; in our case, 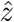 can be thought as a random variable encoding properties associated with enzymes catalyzing a certain reaction.

However, learning the parameters *θ* of this class of models is usually intractable, since we cannot evaluate or differentiate the marginal likelihood ∫ *p*_*θ*_(*x*|*z*)*p*_*θ*_(*z*) and the posterior probability *p*_*θ*_(*x*|*z*)*p*_*θ*_(*z*)/*p*_*θ*_(*x*). Here we addressed these issues by using a Variational Autoencoder (VAE) architecture, where Neural Networks (NNs) are used to approximate *p*_*θ*_(*x*|*z*) and *p*_*θ*_(*z*|*x*), and Stochastic Variational Inference (SVI) to learn the corresponding parameters [25]. Specifically, in a VAE framework, *p*_*θ*_(*z*|*x*) is approximated by a parametric recognition model, *q*_*ϕ*_(*z*|*x*), which act as a probabilistic encoder taking in input a sequence *x* and returning a distribution over the possible value of *z*. Conversely, in this framework, *p*_*θ*_(*x*|*z*) act as a probabilistic parametric decoder, which takes in input a sample *z* and returns a distribution over the possible values of *x*.

Hereby, we introduce the parametric distributions used to model the latent space, the NNs used for encoding and decoding biological sequences, and the optimization procedure used to fit our models.

### Parametric distributions for latent space sequence modelling

We argued that the ability of VAEs to effectively sample the protein design space and generate new functional variant depends on the parametric family used to model the latent space.

VAEs have traditionally used a multi-variate Gaussian distribution, which has desireable mathematical properties (e.g closed analytical forms for the gradient), which in turns makes parameters’ estimation efficient. However, protein sequences are mostly characterized by discrete properties (e.g. family membership, species specificity), and sequences themselves are discrete mathematical entities.

Thus, here we explored the use of the Dirichlet distribution as an alternative parametric distribution to model the enzyme design space; this distribution has been routinely used in statistical sequence analysis for many applications, including sequence clustering [26]. From a sequence design perspective, a Dirichlet distribution has the desirable property to efficiently capture data multi-modalities, which is unfeasible with a Gaussian distribution [27], and thus theoretically superior to model the vast multi modal enzyme design space. To test experimentally this hypothesis, we contrasted and compared sequences generated by VAEs using both the classical multi-variate Gaussian and the Dirichlet distribution.

### Efficient encoding and decoding of biological sequences

A plethora of NNs have been proposed to model sequence data, including Recurrent Neural Network (RNN)[28], Long Short Term Memory (LSTM)[29] and Gated Recurrent Unit (GRU)[30]; however, as the length of the sequences increases, their ability to learn long-range relationships between amino acids decreases [31], a drawback that makes them them unsuitable to handle long amino acid sequences. Moreover, these architectures are computationally expensive to train [32], as they cannot be readily parallelized, and thus unsuitable to scale over large sequence datasets.

Therefore, we used an alternative architecture for both the encoder and decoder, called Temporal Convolutional Network (TCN), which overcomes these limitations and can be efficiently trained [33]. TCNs take sequences in input and return new ones of the same length, which are obtained by sampling from a learned language model. Sequences of the same length are easily obtained by using a standard 1-dimensional convolutional (1D-CONV) layer, using zero-padding to keep the output of the subsequent layers equal to the input length. To condition the probability of a residue on the previously observed ones, TCNs use causal convolution, where the residue at position *t* is obtained by applying convolution only with elements at positions *t*, …, 0 in the previous layers. However, obtaining an effective memory usually requires stacking multiple convolutional layers, thus making vanilla TCNs inefficient to train. The problem has been recently address by using dilated convolution. Let *x* be a sequence of length *n* and *f* a kernel of size *k*, the dilated convolution *F* is computed as:

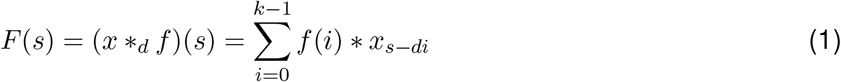

where *d* is the dilation factor, and represent the distance between two elements of the input that are used to produce one element of the output. Stacking multiple temporal convolution layers with exponentially increased dilation factors, allows to obtain full sequence coverage while keeping the number of layers logarithmic in sequence length.

### Variational inference of model parameters

In a Variational Autoencoder framework, NNs are used to compute the variational parameters *ϕ* for a fixed family of probability distributions *q*_*ϕ*_(*z*|*x*) and model parameters *θ* for conditional likelihood *p*_*θ*_(*x*|*z*). Here, we use SVI to find an approximate solution to the problem of maximizing the marginal likelihood *∫p*_*θ*_(*x*|*z*)*p*_*θ*_(*z*) by maximizing the Evidence Lower BOund (ELBO) w.r.t both model parameters *θ* and variational parameters *ϕ* as follows:

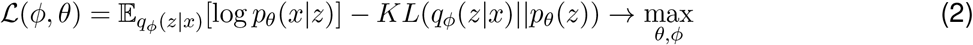

where KL is the Kullback-Liebler divergence, and the expected conditional likelihood 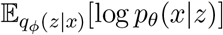 is optimized with respect to maximum log-likelihood, as the probability distribution *p*_*θ*_(*x*|*z*) over the amino acid space is categorical.

Depending on the choice of parametric families for *q*_*ϕ*_(*z*|*x*) and *p*_*θ*_(*x*|*z*) computing the expected conditional likelihood can be challenging, is often intractable and requiring Monte Carlo (MC) approximation, whereas KL divergence can be computed analytically. Ultimately, we need to compute a gradient of expected conditional likelihood w.r.t parameters 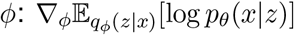. However, in this case, the gradient computation cannot be moved under the expectation operator because the expectation is done w.r.t *q*_*ϕ*_(*z*|*x*), so we revert to the so called reparametization trick [25] to overcome this problem, as follows:

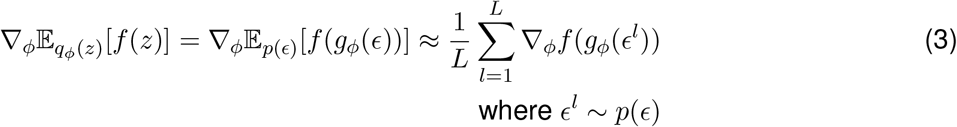

where *p*(ϵ) is a distribution with no parameters to optimize and *g*_*ϕ*_(·) is a deterministic transforma tion of the random variable into another *z*.

For many distributions, the re-parametrization trick is not applicable because no differentiable function *g*_*ϕ*_(·) is available and other approaches, often involving numerical methods, are required. The implicit reparametrization gradient, which can handle the vast majority of distributions and can be applied to any distribution with numerically tractable Cumulative Density Function (CDF), can be introduced as follows:

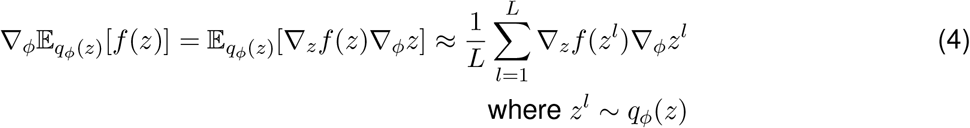

where the gradient 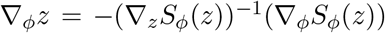, and *S*_*ϕ*_(*z*) is called a standardization function, that is a function that transforms a sample from *q*_*ϕ*_(*z*) into a parameter-free sample. A CDF can serve as a standardization function because it transforms samples from any distribution into standard uniform samples. Thus, in a univariate case, we can write 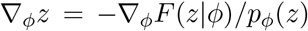, where *F* (·) and *p*(·) are CDF and Probability Density Function (PDF) of a distribution. However, the two approaches differ in the multivariate case, where either a multivariate distributional transform is used [34] or reparametrization gradients are viewed as solutions to a differential equation [35].

### Implementation

Our VAE architecture uses TCNs both as encoder and decoder, and consisting of stacked residual blocks with exponentially increased dilations. Each residual block consists of a pair of temporal convolutional layers followed by weight normalization and dropout and a residual connection [33]. For temporal convolutions, we use intermediate layers of 256 hidden units, dilation factor *d* = 2, kernel size *k* = 32, and subject to 20% dropout during training. The total number of stacked layers was determined automatically as follows:

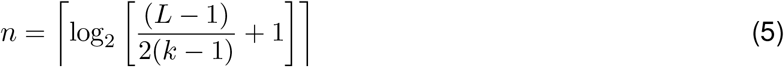

where *L* is the length of the longest sequence. For representing amino acids and special tokens denoting the start and the end of a sequence, we used a 32-dimensional embedding for amino acids subject to a 20% dropout during training.

We tested this architecture using both a Gaussian and Dirichlet latent distribution, which will be hereby denoted as Temporal Gaussian Variational Autoencoder (TGVAE) and Temporal Dirichlet Variational Autoencoder (TDVAE), respectively.

## Results

We tested and evaluated our TGVAE and TDVAE models by designing a new set of human Sphyngosine-1-phosphate lyase (HsS1PL) enzymes, a 568 residues enzyme critical for the physiological function of the sphingolipid pathway. To do that, we first built a dataset of S1PL enzyme and corresponding orthologous sequences in eukaryota from EggNog (v5.0.0, download date: 26/10/2021, [36]). We then removed duplicates, sequences with non-canonical amino acids, and those shorter than 200 and longer than 600 residues. Taken together, our dataset consists of 1,147 sequences with an average length of 493 amino acids. We then performed sequence clustering using MMSEQS2 [37], with minimum sequence identity set to 0.7, using cluster representatives as sequences for the validation dataset and the remaining one as sequences for the training set, ensuring a 90/10 ratio of sequences between training and validation sets by reallocating randomly selected sequences.

We then tested our models by using a 64 and 128 dimensional latent space, while using the same TCN configuration for both the encoder and decoder layers. For each configuration, we performed 10 independent runs, consisting of 99,000 batch updates each, and selected the model parameters associated with the best validation ELBO.

We found that TGVAE achieved the best ELBO on average compared to TDVAE (see Tab. 1), whereas a 64 dimensional latent space always led to lower ELBO regardless of the model used. We then inspected the contribution of the reconstruction error and KL divergence term; we found that TGVAE achieved a lower reconstruction error compared to TDVAE, albeit this result was also associated with higher performance variability. When we looked at the KL divergence, we found the term to be significantly greater than zero for all models, confirming that our models effectively uses the latent space to encode sequence information [38].

**Table 1:**
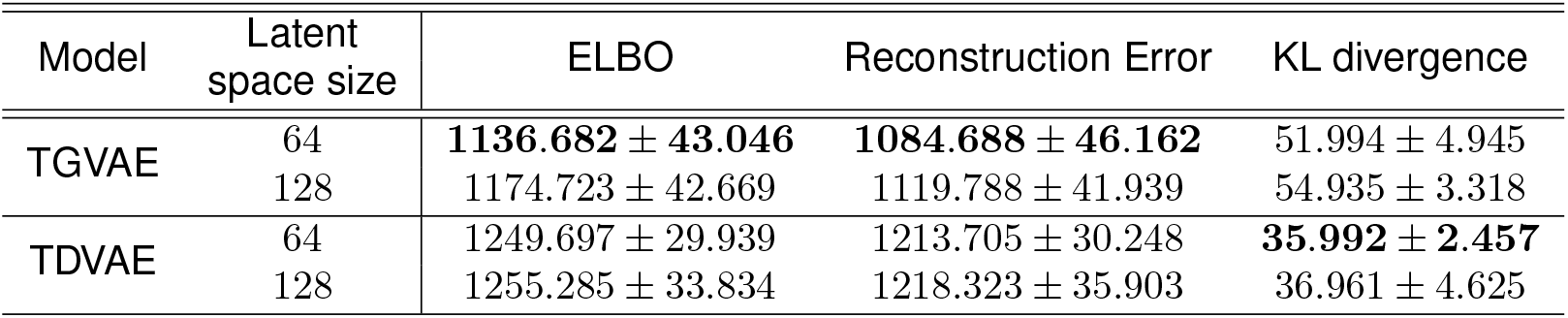
Evaluation of training results for the Temporal Gaussian Variational Autoencoder (TGVAE) and the Temporal Dirichlet Variational Autoencoder (TDVAE). For each model, we report the size of the latent space, the mean and standard deviation of the ELBO over 10 runs, along with the mean and standard deviation of the reconstruction error and KL divergence. In bold, we report the best value for each metric.

Taken together, we found that TGVAE leads to slightly better training performances. However, generative models should be primarily evaluated based on the ability to produce new sequences that resemble the primary and tertiary structure properties on a given enzyme class. Thus, we proceeded to perform a sequence and structural analysis of the variants generated by our models, albeit restricting our experiments to a 64-dimensional space as it was shown to achieve the best performances for both models.

### Sequence analysis of new sphyngosine-1-phosphate lyase enzymes

We then evaluated the ability of our models to design new putatively functional S1PL variants, using an extensive set of sequence similarity metrics and by analyzing their biochemical properties.

To do that, we used two different design strategies, which differ in whether *z* is sampled from the prior distribution (*N* (0, I) for TGVAE and *Dir*(*α* = 1) for TDVAE) or a posterior distribution, whose parameters where obtained by processing an input seed sequence through the encoder; in our case, sampling from the prior is somewhat similar to generating S1PL enzymes de-novo, whereas sampling from the posterior is similar to designing variants of a known enzyme.

First, we trained a 64-dimensional TGVAE and TDVAE and selected the parameters leading to the best validation ELBO. We then generated 100, 000 sequences from the respective prior distributions, and from the posterior distributions obtained by seeding the model with the human Sphyngosine-1-phosphate lyase (HsS1PL) sequence (Uniprot id: O95470). We then computed the number of unique samples generated, the average identity, similarity and bit score of the sequence ensemble as well as 90% empirical confidence interval; specifically, we compared sequences sampled from the prior against the entire S1PL training set and S1PL seeded sequences against HsS1PL, using BLASTP and considering only sequences with E-value < 10^−4^.

We first analysed sequences generated from the respective prior distributions for TGVAE and TDVAE (see Tab. 2); here, we found that TDVAE generated a larger set of unique samples (98.81%) compared to TGVAE (95.55%). When we analysed the sequence similarity metrics, we found the Dirichlet model to generate sequences that are significantly more similar on average to known S1PL enzymes compared to those generated when using a Gaussian model (see Tab. 2). Nonetheless, the low average identity and similarity scores suggests that sequences sampled from the prior are consistently different from known S1PL sequences; while this is a desireable property, as it increases variants diversity, it obviously comes at the expense of an higher rate of putative non-functional enzymes.

**Table 2:**
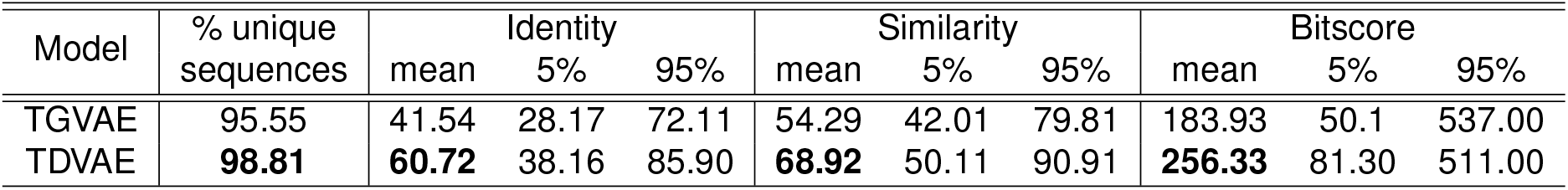
Sequence analysis of variants generated by sampling from the respective prior distributions for TGVAE and TDVAE. For each model, we report the percentage of unique variants from an ensemble of 10^5^ generated sequences, and the corresponding average and 90% empirical confidence interval of identity, similarity and bitscore with respect to S1PL sequences in the training set. In bold, we report the best value for each metric.

| Model | % unique sequences | Identity |  |  | Similarity |  |  | Bitscore |  |  |
| --- | --- | --- | --- | --- | --- | --- | --- | --- | --- | --- |
|  |  | mean | 5% | 95% | mean | 5% | 95% | mean | 5% | 95% |
| TGVAE | 95.55 | 41.54 | 28.17 | 72.11 | 54.29 | 42.01 | 79.81 | 183.93 | 50.1 | 537.00 |
| TDVAE | **98.81** | **60.72** | 38.16 | 85.90 | **68.92** | 50.11 | 90.91 | **256.33** | 81.30 | 511.00 |

We then looked at the sequences obtained by sampling from posterior distribution of HsS1PL. Here, both models generated sequences that are significantly more conserved than those generated by sampling from the prior distributions, with the TDVAE achieving the best results on average (see Tab. 3). Surprisingly, despite all models producing sequences with similarity and identity > 90%, they still provided a set of almost 100% unique sequences, suggesting that the model is learning to sample unseen variants in the neighborhood of the HsS1PL sequence space.

**Table 3:**
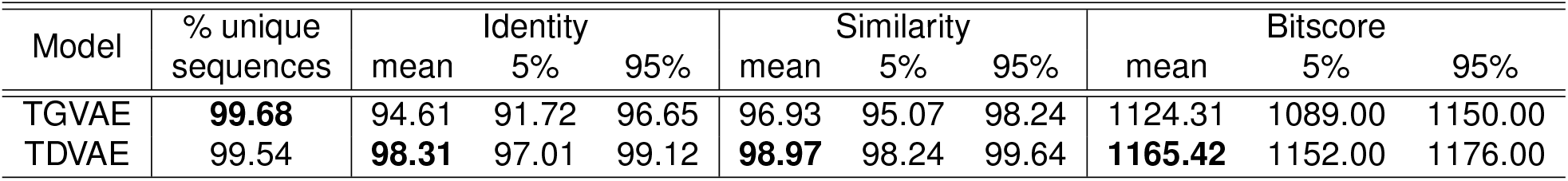
Sequence analysis of variants generated by sampling from the respective posterior distributions for TGVAE and TDVAE. For each model, we report the percentage of unique variants from an ensemble of 10^5^ generated sequences, and the corresponding average and 90% empirical confidence interval of identity, similarity and bitscore with respect to the HsS1PL sequence. In bold, we report the best value for each metric.

| Model | % unique sequences | Identity |  |  | Similarity |  |  | Bitscore |  |  |
| --- | --- | --- | --- | --- | --- | --- | --- | --- | --- | --- |
|  |  | mean | 5% | 95% | mean | 5% | 95% | mean | 5% | 95% |
| TGVAE | **99.68** | 94.61 | 91.72 | 96.65 | 96.93 | 95.07 | 98.24 | 1124.31 | 1089.00 | 1150.00 |
| TDVAE | 99.54 | **98.31** | 97.01 | 99.12 | **98.97** | 98.24 | 99.64 | **1165.42** | 1152.00 | 1176.00 |

As TDVAE generated the largest number of distinct near wild type HsS1PL variants, we further analysed these sequences by comparing their biochemical properties with those of HsS1PL; here, we focused on the protein grand average of hydropathicity index (GRAVY) [39], instability index [40] and isoelectric point [41]. The GRAVY metric allows us to estimate whether a protein is hydrophobic (positive scores) or hydrophilic (negative scores). The instability index, instead, predicts protein instability as a function of the occurrence of destabilizing dipeptides in the sequence, where values greater than 40 are usually indicative of unstable proteins. Finally, as a proxy to estimate protein solubility, we calculated the isoelectric point of each sequence. Here we found that sequences generated from the posterior distribution for HsS1PL have biochemical properties comparable to the wild type enzyme (see Fig. 1), including same hydrophilic propensity, with an average GRAVY of −0.10 compared to −0.11 in wild type, and same isoelectric point, with an average index of 9.24 compared to 9.28 for the wild type. Importantly, sequences from the posterior are predicted to be highly stable with an average index of 32.31, compared to 32.00 of HsS1PL. Notably, when we carried out the same analysis for sequences generated from the prior distribution, we observed a large fraction of proteins being predicted as unstable and hydrophobic.

**Figure 1:**
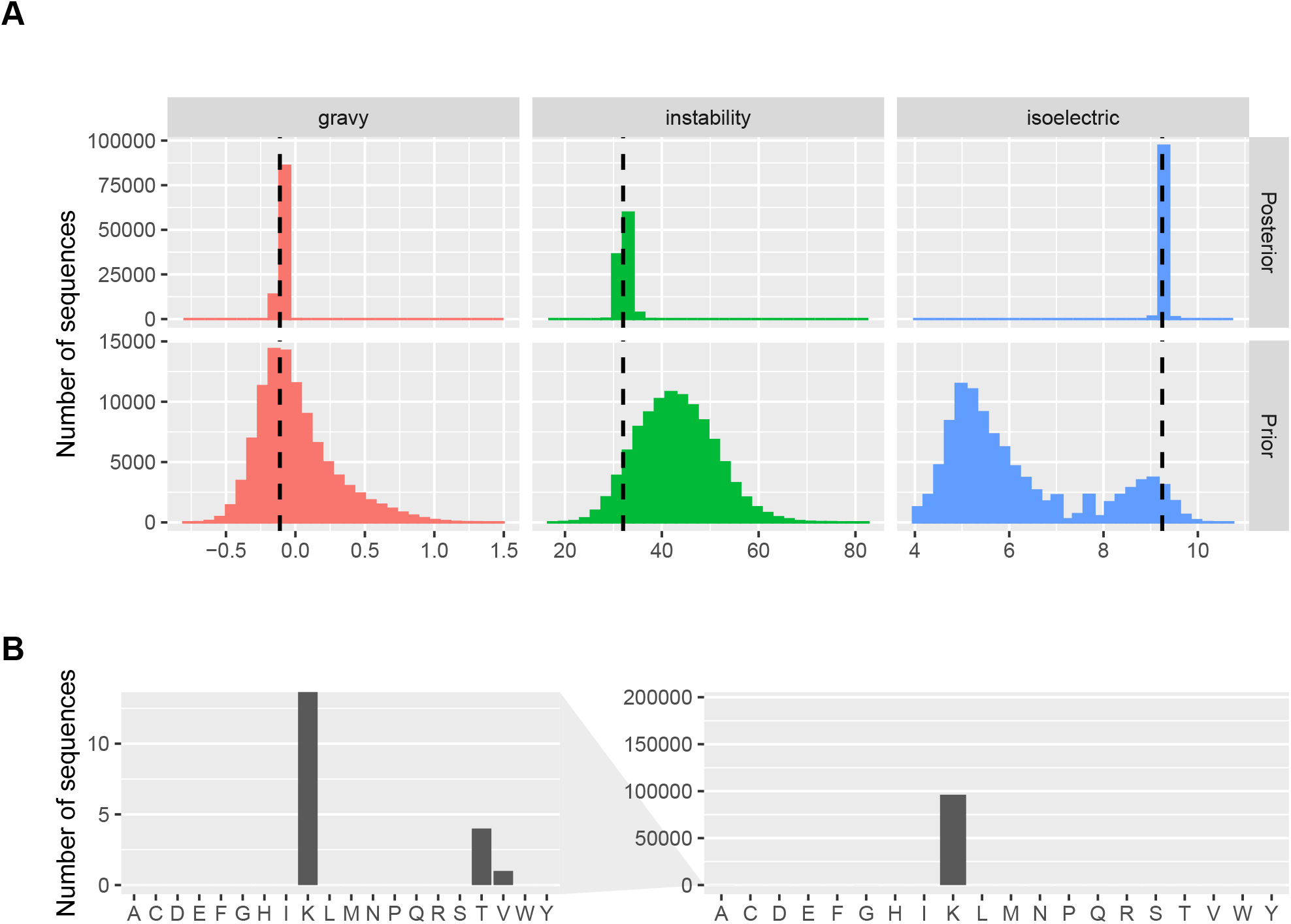
Biochemical properties of HsS1PL variants generated by TDVAE. **A)** Distribution of the flexibility, instability and isoelectric point of the sequences generated from the variational posterior distribution identified by HsS1PL and the prior distribution. While variants generated from the posterior strongly retain wild type properties (dashed line), sequences generated from the prior distribution are considerably more unstable and less soluble. **B)** Number of variants mutating the K_353_ active site in HsS1PL; only 5 variants introduced mutations by replacing lysine with either a threonine or a valine (left side zoom plot).

Finally, we looked whether variants generated by TDVAE retained the active site *K*_353_ (see Fig. 1). Here we found the activate site to be conserved across all variants but 5, where it was substituted by a threonine (4 variants) and a valine(1 variant).

Taken together, we found that TDVAE generated a large and diverse set of sequences with chemical properties similar to HsS1PL, thus providing a potentially large set of functional HsS1PL variants.

### Structural analysis of human Sphyngosine-1-phosphate lyase variants

We have shown that TDVAE can generate HsS1PL variants that retain the biochemical properties of the wild type sequence, while adding up to 50 mutations across the sequence. However, sequencebased assessment of enzyme function provides only limited insights on the designed variants, and prioritization strategies based on sequence alone would neglect structural properties, such as ligand docking, that cannot be captured by sequence-based assessment methods. Thus, we set up an extensive structural analysis protocol to: i) understand where mutations are located with respect to the wild type structure, and ii) assess the stability and structural integrity of the HsS1PL variants.

To investigate sequence variability across variants, we selected the top 200 sequences ranked by log-likelihood, performed multiple sequence alignment using Clustal Omega and calculated entropy-based conservation scores at each position using AL2CO [42]. These conservation scores were then mapped onto the wild type structure of HsS1PL (PDB id: 4Q6R) for ease of visualization (see Fig. 2). Here, we found that amino acid changes are mostly located on or nearby coiled regions; importantly, we found the regions defining the PLP binding site – including the critical K_353_ and pocket residues G_210_, T_211_, H_242_, C_317_, L_318_, G_394_, Y_387_, G_394_ – to be completely conserved, which suggests that TDVAE can preserve the critical functional elements of the enzyme.

**Figure 2:**
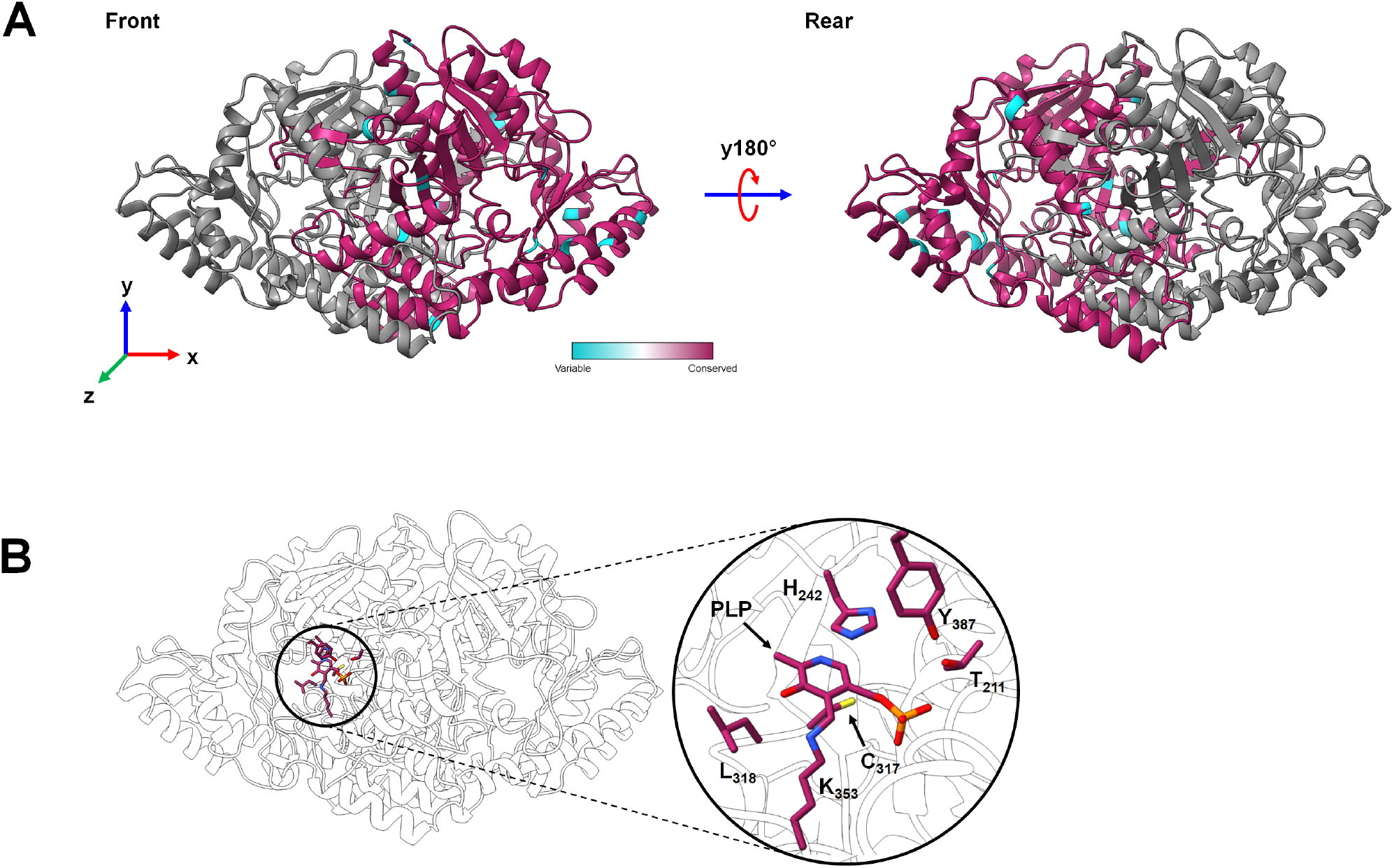
Structural analysis of mutations in HsS1PL variants. A visualization of sequence conservation using the HsS1PL crystal structure (PDB id: 4Q6R) derived from the multiple sequence alignment of the top 200 variants generated by TDVAE. **A**) Entropy-based conservation scores mapped onto chain B of the HsS1PL, highlighting positions mutated by the TDVAE model. Chain A is coloured in grey. **B**) A magnified view of some highly conserved pocket residues that bind and stabilise the internal aldamine formed between the PLP cofactor and the catalytic lysine (K_353_).

Then, to estimate the impact of the mutations on protein structure and stability, we randomly selected 7 variants sufficiently different from the set of sequences with identity between 90% − 100% from the wild type, and which were predicted to be stable but still carrying a significant number of mutations. For each sequence, we then predicted their structure using COLABFOLD [43, 44], configured to build multiple sequence alignments (MSA) using MMSEQS2, to perform homomeric prediction without templating, and allowing 3 rounds of structural recycling; we then selected the best model ranked by pTM for downstream molecular dynamic (MD) simulations (see Tab. 4). Importantly, the accurate prediction of the dimer interface is critical for the proper definition of the active site, as it requires residue contributions from both monomeric chains. Finally, as a positive control, the structure of the wild type sequence was also predicted using the aforementioned parameters.

**Table 4:** Structural analysis of HsS1PL. For each variant, we report a mnemonic sequence identifier (SID), the percent identity with respect to HsS1PL, the ALPHAFOLD2 model confidence score (pLDDT) and structural alignment metric (pTM), along with the RMSD with respect to the core region of the enzyme (E_111_-D_553_). We also evaluated the quality of the predicted interface against HsS1PL by reporting the *F_nat_*, iRMS, LRMS and DOCKQ metrics. For comparison, we also report the metrics computed on the predicted structure for the wild type HsS1PL sequence (O95470). In bold, we report the best value of each metric.

| SID | Identity | pLDDT | pTM | RMSD<br>(Pruned) | RMSD<br>(Global) | $F_{nat}$ | iRMS | LRMS | DockQ |
| --- | --- | --- | --- | --- | --- | --- | --- | --- | --- |
| O95470 | - | 96.170 | 0.949 | 0.439 | 0.545 | 0.930 | 0.589 | 0.788 | 0.929 |
| D4 | **99.648** | **96.040** | **0.950** | 0.467 | 0.565 | 0.924 | 0.607 | **0.846** | 0.925 |
| D1008 | 99.472 | 96.020 | 0.950 | 0.471 | 0.565 | 0.943 | 0.598 | 0.837 | 0.932 |
| D39595 | 97.887 | 96.020 | 0.949 | 0.491 | 0.573 | 0.926 | 0.602 | 0.827 | 0.926 |
| D47945 | 97.535 | 96.020 | 0.949 | 0.480 | 0.573 | **0.945** | 0.611 | 0.828 | 0.931 |
| D90903 | 96.303 | 95.940 | 0.949 | 0.451 | 0.544 | 0.922 | 0.590 | 0.798 | 0.927 |
| D96413 | 95.070 | 95.910 | 0.949 | **0.427** | **0.534** | 0.939 | 0.569 | 0.758 | **0.935** |
| D97533 | 93.662 | 95.700 | 0.946 | 0.482 | 0.575 | 0.935 | **0.612** | 0.781 | 0.928 |

We obtained high-confidence predictions for all variants with pLDDT> 90% and pTM> 0.9. All models, including the wild type, exhibit strong similarity to the solved wild type structure (global RMSD vs. 4Q6R < 0.6AÅ), despite the absence of template modelling. An assessment of the homodimeric interface was performed using DOCKQ[45], which evaluates the quality of a predicted interface against a specified native structure (typically an experimentally-solved structure). DOCKQ computes three metrics proposed and standardized by the Critical Assessment of PRedicted Interactions (CAPRI) community, namely the fraction of preserved native contacts (*F*_*nat*_), the interface Root Means Square Deviation (iRMS) and the Ligand Root Means Square Deviation (LRMS) [46]. DOCKQ combines these metrics into a single score in the [0, 1] range; a DOCKQ score > 0.8 suggests that the predicted interface is highly accurate. Using the native structure reference (PDB id: 4Q6R), we found that variants preserved almost all the native interfacial contacts (*F*_*nat*_ > 0.92), with all complexes returning a DOCKQ score > 0.9. Together with the pTM score, which provides a measure of confidence in the relative orientations of the subunits in a multimeric model, we concluded that the predicted complexes were of excellent quality and suitable for subsequent molecular dynamics (MD) simulation.

We then performed MD simulations to study protein stability over time, using GROMACS 2021.4 [47] with the CHARMM36 all-atom force field [48]. Protein models were solvated in TIP3P water in a cubic box and the net protein charge was counterbalanced using chloride ions. The system underwent potential energy minimization by gradient descent and equilibrated to 300K and 1 bar using V-Rescale thermostat/Berendsen barostat. Following a 10 ns (5 × 10^6^ time steps) production MD, trajectories were re-centered around the catalytic lysine residues (K_353_) with additional rotational and translational fitting. PDB: 4Q6R was also simulated as a positive control. Three attributes were evaluated for each trajectory, namely the average radius of gyration, the pairwise RMSD (2D RMSD) and the inter-chain hydrogen bonding (see Tab. 5).

**Table 5:**
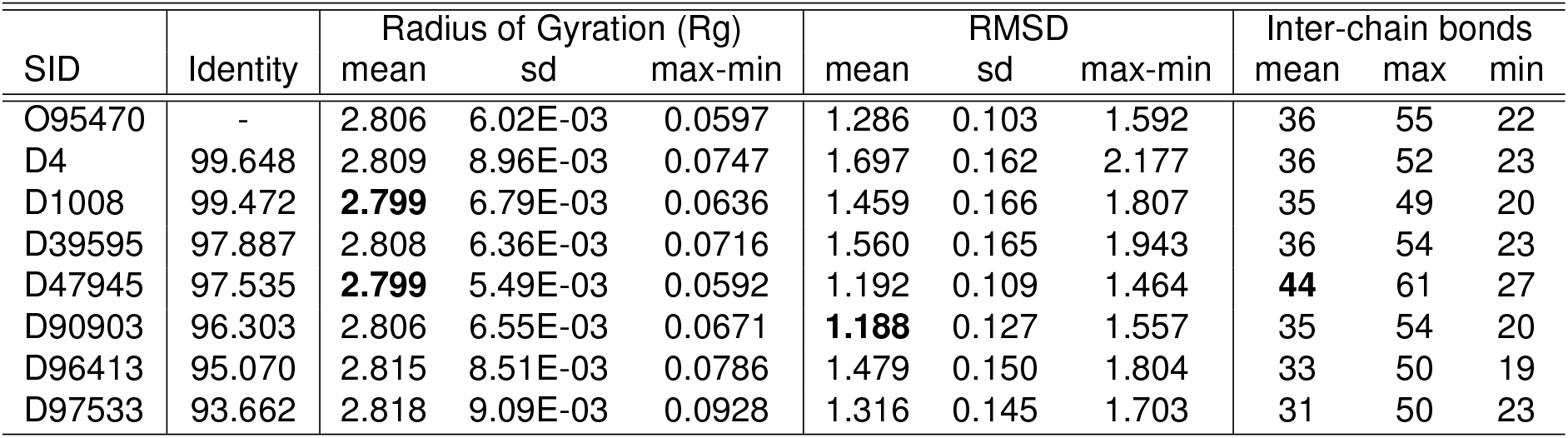
Molecular dynamics analysis of HsS1PL variants. For each structure, we report the associated mnemonic sequence identifier (SID), the percent identity respect to the wild type sequence, the Radius of Gyration (Rg), the RMSD of the structures at each time with respect to the structure at time *t* = 0ns, and the number of inter-chain hydrogen-bond contacts between the chains in the HsS1PL homodimer. For comparison, we also report the metrics computed on the predicted structure for the wild type HsS1PL sequence (O95470). In bold, we report the best value of each metric.

The average Rg for the simulated HsS1PL complexes ranges from 2.80 − 2.82*nm*, while the difference between the maximum and minimum gyrational radii (Rg max-min) averages less than 0.1*nm*. Furthermore, each variant model is stabilized by a similar number of inter-chain hydrogen bonds to the wild type structure, with a notable exception being D47945 which exhibits an average 44 interchain hydrogen bonds over the course of the trajectory. D47945 also returns one of the smallest average Rg (2.799 ± 5.49^−3^*nm*) and the narrowest Rg max-min (0.0592*nm*), suggesting that this complex is particularly compact. Across all variant models, the distance distribution of inter-chain H-bonds is remarkably similar to the simulated wild type structure, with the majority (52.7%) of H-bonds occurring within a distance of 2.725 − 2.925AÅ. Taken altogether, our analysis suggests that the variant S1PL sequences form highly stable homodimeric complexes that maintain their integrity.

To obtain a more qualitative assessment of structure stability over time, we computed pair-wise RMSD in UCSF Chimera [49] between structures at each time step (see Fig. 3). Surprisingly, we observed significant differences between variants, despite sequence identity being greater than 90% for all sequences. In particular, variant D4 exhibits the largest C_*α*_ deviations despite bearing the highest sequence identity to the wild type (99.65%). In contrast, sequences D47945 (97.54%) and D90903 (96.30%) remain comparatively rigid (self-similar) over the course of their respective trajectories. It is interesting to note, however, that all sequences display some degree of heightened flexibility compared to the 4Q6R control. To identify the source of this observed structural mobility, per-residue B-factors were calculated for each trajectory and mapped onto their respective energy-minimized structures for ease of visualization (see Fig. 4). In all instances, the most mobile regions are located on the periphery of the structure, situated on or around coils. Importantly, the core of the proteins remain resilient to fluctuation, as evidenced by the comparatively smaller B-factors of structured and solvent-buried regions.

**Figure 3:**
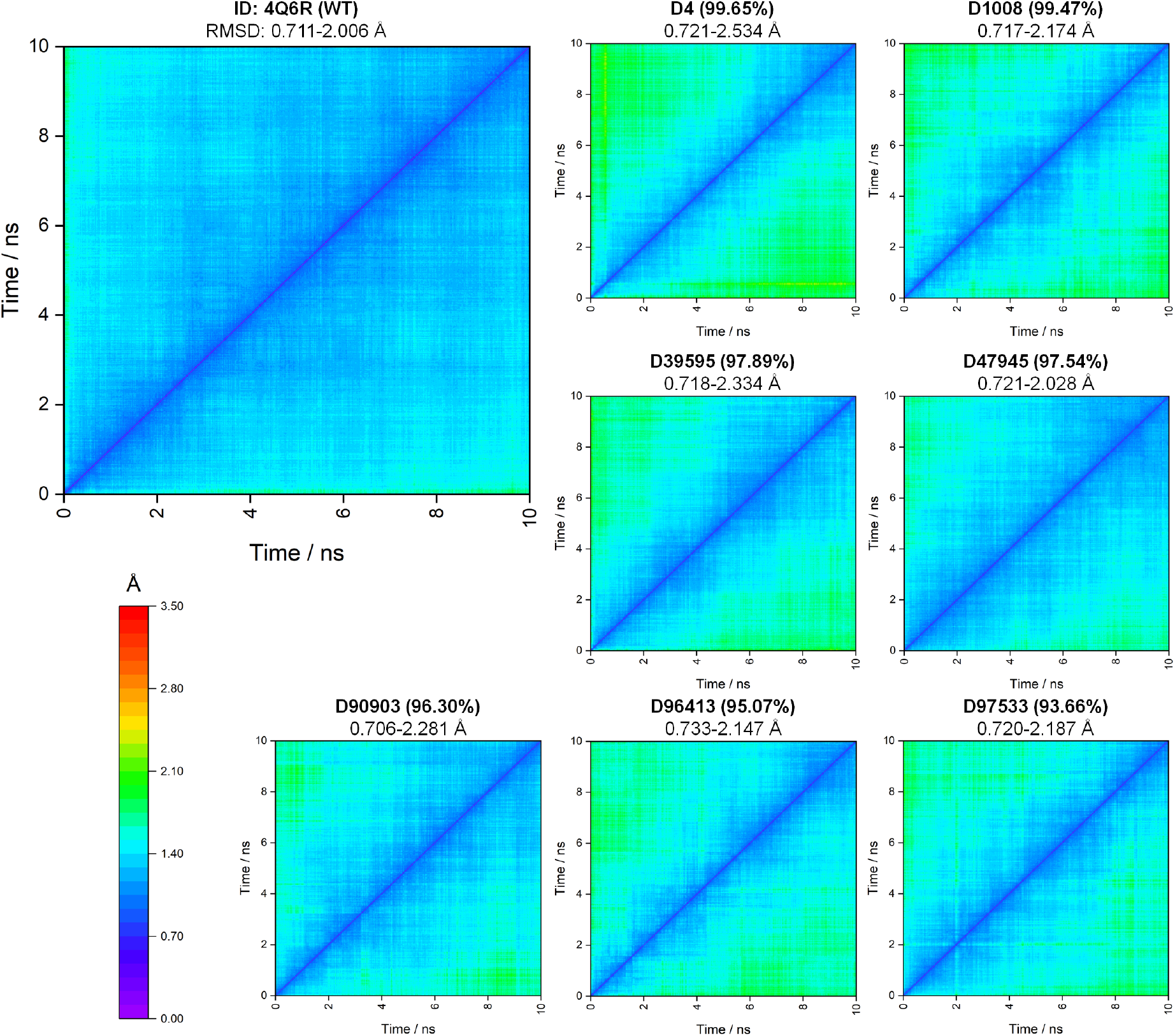
2D RMSD analysis of HsS1PL variants. For each variant, we report a heatmap of the pairwise RMSD of the structures obtained at each time step of the corresponding 10ns molecular dynamics simulation.

**Figure 4:**
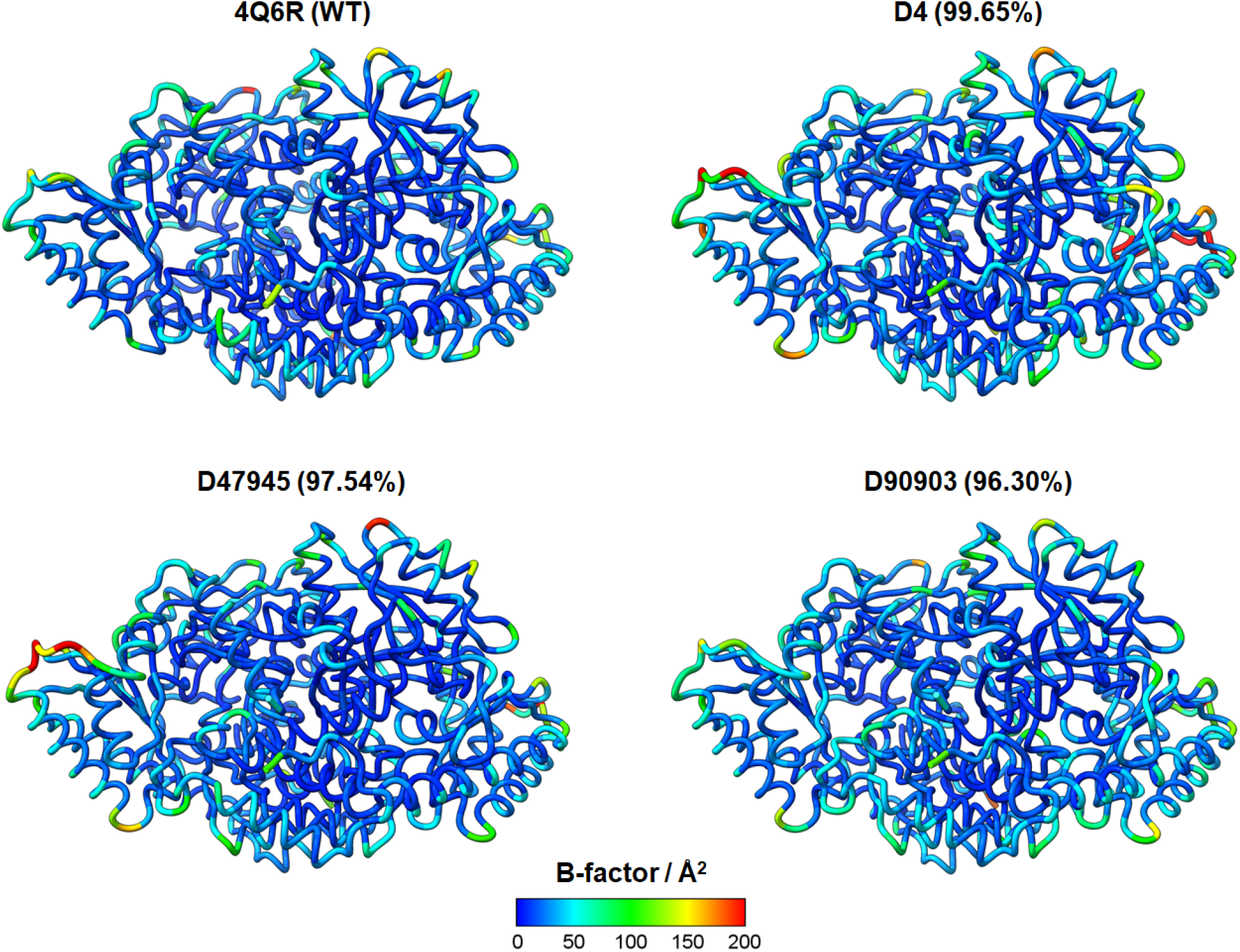
B-factors analysis of HsS1PL variants. Visualization of per-residue B-factors computed for models 4Q6R, D4, D47945 and D90903, computed from their respective trajectories. Larger B-factors are indicative of greater mobility.

It is not yet possible to predict how these fluctuations will impact biochemical fitness; however, our results suggest that the functional landscape of HsS1PL is expected to be rugged, and even single mutations in critical point could lead to non-functional variants, regardless of their sequence similarity with respect to wild type.

## Discussion

Enzyme replacement therapies are the standard of care to treat rare enzymatic deficiencies, which consist in the injection of a recombinant enzyme to restore physiological metabolic activity and improve patients’ symptoms. However, engineering recombinant enzymes with desirable therapeutic properties has been challenging.

Here we showed how deep generative models can be exploited to design variants of human enzymes with structural and biochemical properties similar to known functional wild type molecules, thus providing a tool to expand the enzyme repertoire available for downstream therapeutic applications. Specifically, we developed a new Variational Autoencoder (VAE), called Temporal Dirichlet Variational Autoencoder (TDVAE), which combines a highly efficient Temporal Convolutional Networks (TCNs) for encoding and decoding sequences with Stochastic Variational Inference (SVI) for fast parameters learning, while using a Dirichlet distribution to model the enzyme design space.

As a proof of concept, we used TDVAE to design variants of the human Sphyngosine-1-phosphate lyase (HsS1PL) enzyme, as potential therapeutic enzymes to treat nephrotic syndrome type 14 (NPHS14). Experimental results showed that TDVAE generated a large ensemble of HsS1PL variants with preserved functional features, including the presence of the key catalytic lysine residue. Surprisingly, obtaining these variants did not require training on large sequence datasets or multiple sequence alignment information [21], which are usually difficult to obtain when sequences are highly divergent as S1PL. We then further validated our results by predicting the structure of a subset of variants and performing molecular dynamics simulations to assess enzyme structural stability and integrity; here we found HsS1PL variants to maintain favorable inter-chain contacts to form stable, compact and largely invariant homodimeric complexes. Taken together, our generative design strategy identified high-confidence HsS1PL variants for downstream experimental validation.

While we have shown the effectiveness of TDVAE, we also recognize its limitations. TDVAE can generate a large number of wild type like variants, but our design step is only conditioned on the primary structure, which might not be sufficient to capture structural properties. In fact, we have shown, through MD simulations, that selecting candidates based only on the similarity with the wild type sequence might not be sufficient to identify functional enzymes. Thus, it is important to develop models that can capture biological constraints, while allowing sufficient flexibility to build diverse variants libraries for downstream experimental testing. From a mathematical point of view, we also recognize that is still unclear how the latent encoding can be exploited beyond constraining the search space to the local neighborhood of observed enzyme sequences, and provide general design principles to guide enzyme engineering.

Nonetheless, with the increasing number of available protein sequences and structures across the kingdom of life, we expect VAEs to be well powered to provide effective and accessible sequence-to-function models to drive the engineering of the next generation of therapeutic enzymes.

## Contributions

G.S. conceived the study. G.S. and E.L. formulated and developed the model. E.L. implemented and tested the model and performed sequence analysis under G.S. supervision. M.H. performed and analysed molecular dynamics experiments under D.J.C. supervision. G.S. and E.L. wrote the manuscript with contribution from all the authors.

## Acknowledgments

This work was supported by the UKRI EPSRC Fellowship (EP/V033794/1) to G.S and the UKRI Centre for Doctoral Training in Biomedical AI (grant EP/S02431X/1) for E.L. Computational experiments were performed using resources provided by the Cambridge Service for Data Driven Discovery (CSD3) operated by the University of Cambridge Research Computing Service (https://www.csd3.cam.ac.uk), provided by Dell EMC and Intel using Tier-2 funding from the Engineering and Physical Sciences Research Council (capital grant EP/P020259/1), and DiRAC funding from the Science and Technology Facilities Council (https://www.dirac.ac.uk).

## Notes

### Competing Interest Statement

The authors have declared no competing interest.

